## Supplementary Text for "Identifiability of phenotypic adaptation from low-cell-count experiments and a stochastic model"

Supplementary Material for
"Identifiability of phenotypic adaptation from low-cell-count experiments and a stochastic model"

Alexander P Browning[1,2], Rebecca M Crossley[2], Chiara Villa[3,4], Philip K Maini[2], Adrianne L Jenner[5], Tyler Cassidy[6], and Sara Hamis[7]

[1] *School of Mathematics and Statistics, University of Melbourne, Melbourne, Australia*
[2] *Mathematical Institute, University of Oxford, Oxford, United Kingdom*
[3] *Sorbonne Université, CNRS, Université de Paris, Inria, Laboratoire Jacques-Louis Lions UMR, Paris, France*
[4] *Universit Paris-Saclay, Inria, Centre Inria de Saclay, 91120, Palaiseau, France*
[5] *School of Mathematical Sciences, Queensland University of Technology, Brisbane, Australia*
[6] *School of Mathematics, University of Leeds, Leeds, United Kingdom*
[7] *Department of Information Technology, Uppsala University, Uppsala, Sweden*

April 17, 2025

### Contents

**S1 IBM/CME comparison** **2**

**S2 MCMC priors and results** **3**

**S3 CME for discrete model** **4**

**S4 Structural identifiability of heterogeneity** **5**

**S5 Large data set inference with noisy data** **6**

**S6 Inference with correlated data** **8**

#### S1 IBM/CME comparison

In Fig. S1 we compare the probability mass functions arising from the solution of the CME to the empirical distribution arising from 1000 cell proliferation assays simulated using the IBM.

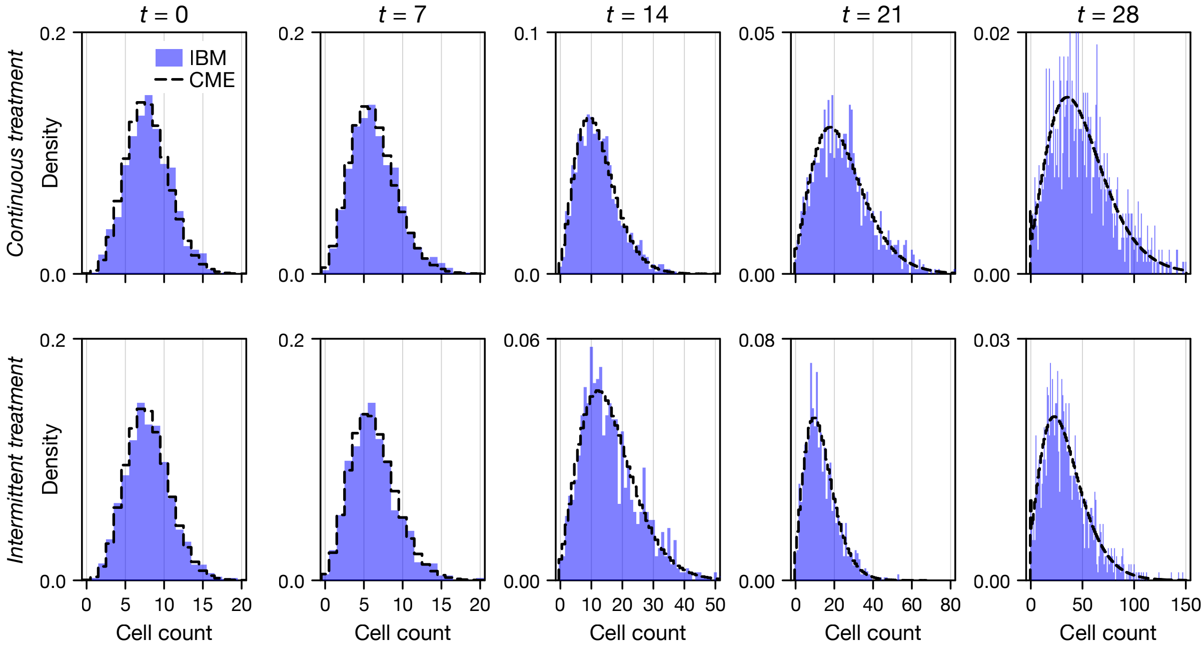


**Figure S1. Chemical master equation comparison.** Comparison between $n = 1000$ realisations of the IBM (blue) and the solution to the CME (black dashed) under continuous treatment (top row) and intermittent treatment (bottom row). All parameters are consistent with those in Fig. 2 of the main text.

### S2 MCMC priors and results

The priors for each parameter are given in Table S1. In Fig. S2, we show marginal posterior distributions for all parameters corresponding to the analysis in Fig. 3 of the main text. In the main text, we present results for experiment termination times $t = \{1\,\mathrm{d}, 3\,\mathrm{d}, 5\,\mathrm{d}, 7\,\mathrm{d}\}$. In Fig. S2, we additionally consider various other termination time sets, as indicated.

**Table S1.** Parameters, parameter descriptions, and parameter priors used for analysis in the main text.

| Parameter | Description | Prior |
|---|---|---|
| $\gamma_1$ | Sensitive $(x = 0)$ growth rate off drug | Uniform$(-1, 1)$ |
| $\gamma_2$ | Sensitive $(x = 0)$ growth rate on drug | Uniform$(-1, 1)$ |
| $\gamma_3$ | Resistant $(x = 1)$ growth rate off drug | Uniform$(-1, 1)$ |
| $\gamma_4$ | Resistant $(x = 1)$ growth rate on drug | Uniform$(-1, 1)$ |
| $\log \nu$ | Adaptation speed | Uniform$(-6, 1)$ |
| $\log \beta$ | Heterogeneity/diffusivity parameter | Uniform$(-6, -1)$ |

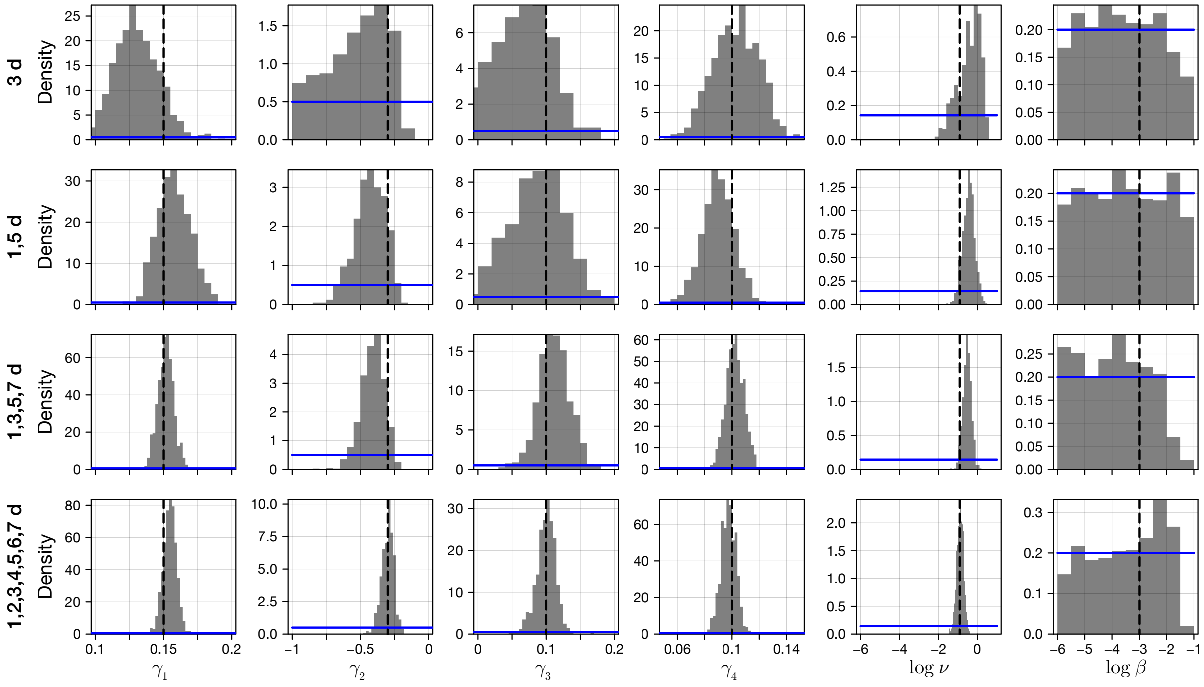


**Figure S2. Marginal posterior distributions.** We show marginal posterior distributions for all parameters, for various sets of observation termination times. Shown also are the marginal prior distributions (blue) and the true values (black dashed).

#### S3 CME for discrete model

Let $q(n_1, n_2, t)$ be the density for $n_1$ sensitive cells and $n_2$ resistant cells. Let $\lambda_i$ denote the proliferation rate of each cell population, and $\delta_i$ denote the death rate of each cell population (one of which will be zero). We have that

$$\begin{aligned}\frac{\mathrm{d}q(n_1,n_2,t)}{\mathrm{d}t} &= (n_1-1)\lambda_1 q(n_1-1,n_2,t) + (n_1+1)\delta_1 q(n_1+1,n_2,t)\\ &\quad + (n_2-1)\lambda_2 q(n_1,n_2-1,t) + (n_2+1)\delta_2 q(n_1,n_2+1,t)\\ &\quad + n_1 r_{12} q(n_1+1,n_2-1,t) + n_2 r_{21} q(n_1-1,n_2+1,t)\\ &\quad - (n_1\lambda_1 + n_1\delta_1 + n_2\lambda_2 + n_2\delta_2) q(n_1,n_2,t).\end{aligned} \tag{1}$$

To obtain the mass function for the total cell count, $q(n,t)$, we consider that

$$q(n,t) = \sum_{n_1=0}^{\infty} q(n_1, n-n_1, t). \tag{2}$$

In practice, we consider a partial sum truncated at $n_1 = 50$, which we find to be sufficient given the maximum cell counts observed in Fig. 8 of the main text.

#### S4 Structural identifiability of heterogeneity

In the main text, we observe that the heterogeneity parameter, $\beta$, is one-sided practically identifiable. In this section, we repeat the computational experiment associated with Fig. 3 of the main text (and equivalently, Fig. S2 of this supplementary material document) to investigate a scenario where a large data set is generated that comprises 768 cell proliferation assays (i.e., eight plates), for each condition, at a set of termination times $t = \{0.5\,\mathrm{d}, 1\,\mathrm{d}, 1.5\,\mathrm{d}, \cdots, 6.5\,\mathrm{d}, 7\,\mathrm{d}\}$. Results in Fig. S3 show that, in this large data-set regime (5,376 proliferation assays equivalent to a total of 56 plates), the diffusivity parameter is practically identifiable, demonstrating that it is a structurally identifiable parameter.

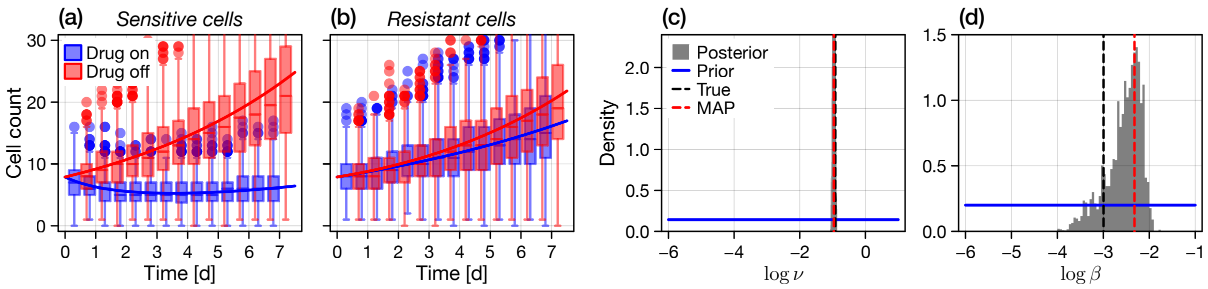


**Figure S3. Large data set proliferation assay inference.** We show marginal posterior distributions for all parameters, for various sets of observation termination times. We perform Bayesian inference on synthetic cell proliferation assay data using the CME as a likelihood. Independent cell count observations ($M = 768$ replicates per condition) are made from experiments conducted with fully sensitive or fully resistant cells, with and without drug, and terminated at $t = \{0.5\,\mathrm{d}, 1\,\mathrm{d}, 1.5\,\mathrm{d}, \cdots, 6.5\,\mathrm{d}, 7\,\mathrm{d}\}$. (a–b) Synthetic proliferation assay cell count data (box plots with outliers shown as discs) and the model predicted mean cell count at the MAP (solid lines). (c–d) Posterior distributions for the logarithm of $v$, the adaptation speed, and $\beta$, the diffusivity. Shown also is the uniform prior (blue), the true value (black dashed), and the MAP (red dashed). In this regime, both the adaptation speed and diffusivity are identifiable.

#### S5 Large data set inference with noisy data

In the main text, and in all supplementary results hitherto, we have studied a scenario where exact cell counts are made. We find in Section S4 that, in this effectively noise-free regime, the heterogenity parameter is identifiable provided that a sufficiently large data set is available. We now revisit this assumption by assuming that cell count observations, denoted $y$, are subject to binomial noise such that

$$y \,|\, n \sim \text{Truncated}\left(\text{Binomial}(M(n), 0.5) - \frac{M(n)}{2} + n, 0, \infty\right), \tag{3}$$

where $n$ is the exact cell count, and $M(n)$ is chosen such that the standard deviation of the noise term scales with the cell count (note that for the right hand side of Eq. (3) to be valid, we require that $M(n)$ is always even, such that the noise term $\text{Binomial}(M(n), 0.5) - M(n)/2$ is symmetric about zero. In this section, we set

$$M(n) = 2 \cdot \text{round}\left(\frac{4\alpha^2 n^2 + n_0}{2}\right), \tag{4}$$

such that the noise comprises a count independent term, $n_0$ (i.e., noise present even in very low cell count observations arising from, e.g., cellular debris), and a count dependent term of magnitude $\alpha$ which scales such that the standard deviation of the noise term is approximately $\alpha n$ for large $n$. For the results that follow, we set $\alpha = 0.1$ and $n_0 = 5$; for these parameter values we demonstrate the noise distribution and resultant observed cell count distribution in Fig. S4. While complex, this choice of discrete noise model accounts for both over and under counting (for example, an automated counting algorithm that both misses cells, and misclassifies cellular debris as cells).

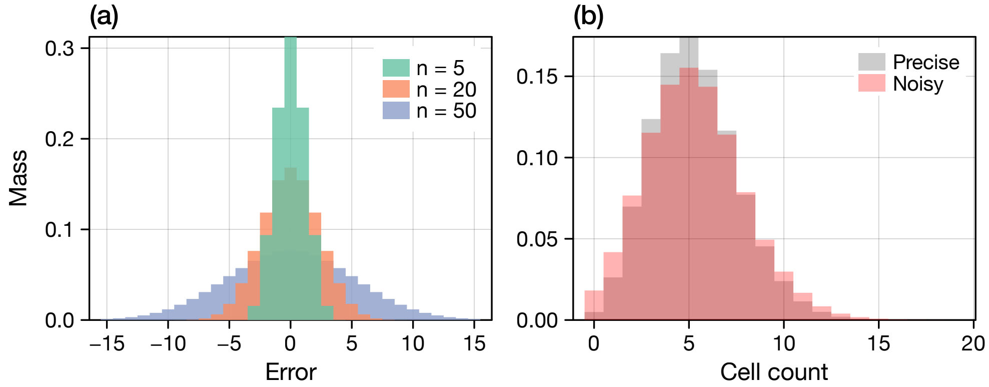


**Figure S4. Observation noise model.** (a) We consider cell count observations subject to additive Binomial error that scales with the cell count, $n$, according to Eq. (4). (b) Comparison between precise (black) and noisy (red) cell count distributions from the CME.

Following the construction of the statistical observation noise model, we reproduce the large data set results of Fig. S3 in the case that only noisy observations are available. Priors are given in Table S2 and both fits and marginal posterior distributions in Fig. S4. Results in Fig. S5 demonstrate that the diffusivity parameter is again only one-sided identifiable; even from a large data set, we cannot distinguish heterogeneity from observation noise.

**Table S2.** Prior distributions for the noise distribution parameters used in the inference of noisy data.

| Parameter | Prior |
|---|---|
| $\alpha$ | Uniform$(0, 1)$ |
| $n_0$ | Uniform$(0, 10)$ |

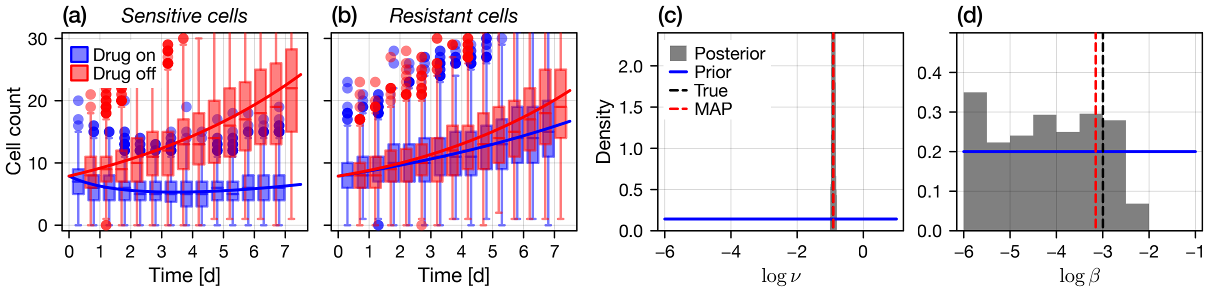


**Figure S5. Large data set proliferation assay inference with noisy data.** We reproduce the results in Fig. S3 in the case that cell count observations are subject to noise of the form given in Eq. (3). The noise parameters $\alpha$ and $n_0$ are assumed to be unknown, with priors given in Table S2.

### S6 Inference with correlated data

For simplicity, in the main document we assume that all cell count observations are independent. For a cell proliferation assay, this is often not true: rather, observations of the *same* cell proliferation assay are made at multiple time points.

In the case that measurements are non-subject to measurement noise, an analogous log-likelihood can be constructed using the chemical master equation. Denote $0 < t_1 < t_2 < \ldots < t_K$ the set of observation times, and $N_1, N_2, \ldots, N_K$ a corresponding set of observed counts. The contribution to the likelihood can be simplified by applying the Markov property of the process, such that

$$\mathbb{P}(N_1, N_2, \ldots, N_K) = \mathbb{P}(N_K|N_{K-1})\mathbb{P}(N_{K-1}|N_{K-2}) \cdots \mathbb{P}(N_1). \tag{5}$$

The term $\mathbb{P}(N_1)$ is simply given by a numerical solution to the chemical master equation at $t = t_1$. Other terms $\mathbb{P}(N_k|N_{k-1})$ can similarly be obtained by the solution to a chemical master equation at $t = t_k$ subject to the initial condition $N(t_{k-1}) = N_{k-1}$.

In Fig. S6, we reproduce Fig. 3 of the main document in the case that $M = 192$ wells are used per condition (a total of two 96-well plates), all observed at the four time points. Thus, the same number of plates are used per condition, but a set of four correlated observations are made of each well. Results in Fig. S6 show that $\beta$ remains one-sided identifiable.

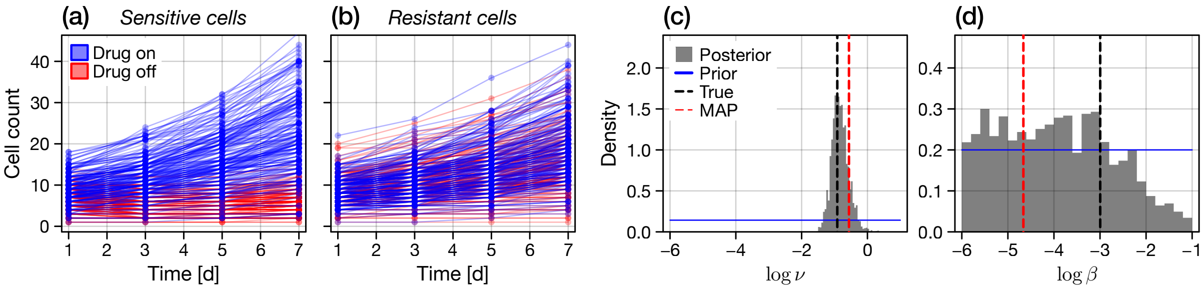


**Figure S6. Inference with correlated data.** We reproduce the results in Fig. 3 of the main document in the case that the data are correlated between observation times.
